# dropClust: Efficient clustering of ultra-large scRNA-seq data

**DOI:** 10.1101/170308

**Authors:** Debajyoti Sinha, Akhilesh Kumar, Himanshu Kumar, Sanghamitra Bandyopadhyay, Debarka Sengupta

## Abstract

Droplet based single cell transcriptomics has recently enabled parallel screening of tens of thousands of single cells. Clustering methods that scale for such high dimensional data without compromising accuracy are scarce. We exploit Locality Sensitive Hashing, an approximate nearest neighbor search technique to develop a *de novo* clustering algorithm for large-scale single cell data. On a number of real datasets, dropClust outperformed the existing best practice methods in terms of execution time, clustering accuracy and detectability of minor cell sub-types.

---

Biological systems harbor substantial heterogeneity, which is hard to decode by profiling population of cells. Over the past few years, technological advances enabled genome wide profiling of RNA, DNA, protein and epigenetic modifications in individual cells^1^. Amongst the most recent developments, in-drop (within a droplet) barcoding has gained a lot of attention as it enables 3′ mRNA counting of thousands of individual cells in a matter of several minutes to few hours. A recent work produced an unprecedented ∼ 250k single cell transcriptomes as part of a single study^2^. This gives us an idea about the scale of the future single cell experiments. Since the introduction of single cell RNA sequencing (scRNA-seq) technologies, a number of clustering techniques have been devised while accounting for the unique characteristics of the new data type^3–6^. However, a majority of these techniques struggle to scale when studies feature several tens of thousands of transcriptomes. In fact, methods developed solely for such ultra large datasets (henceforth referred to as Drop-seq data, following Macosco et al^7^) are either computationally expensive^7^ or over-simplistic^2^, which obscure fast and accurate analysis of such voluminous scRNA-seq datasets.

Network based clustering techniques have been used effectively for clustering single cell transcriptomes^8,9^. An exhaustive nearest neighbor search requires quadratic-time tabulation of pair-wise distances. For large sample sizes, this approach turns out to be significantly slow. Seurat, one of the early-proposed methods for Drop-seq data analysis, performs sub-sampling of transcriptomes prior to nearest-neighbor based network construction, followed by topological clustering. Random sampling can be irreversibly lossy when one of the objectives is to identify rare cell populations. In a recent work, Zheng and colleagues^2^ have used *k*-means as the method for clustering Drop-seq data. Although the choice of *k*-means reduces the overall time for analysis, it suffers from two major drawbacks: 1. User needs to specify the number of clusters. 2. The method struggles to identify clusters of non-spherical shapes.

To address the above shortcomings, we developed dropClust, a scalable yet accurate clustering algorithm for Drop-seq data. DropClust employs Locality Sensitive Hashing (LSH), a logarithmic-time algorithm to determine approximate neighborhood for individual transcriptomes. An approximate *k* nearest neighbor network of individual transcriptomes thus obtained, is subjected to Louvian^10^, a widely used network partitioning algorithm (**Online Methods**). We observed that usually a majority of the distinct subtypes, including many of the rare cell clusters, get identified at this stage. While topological clustering is quite robust in identifying high level subpopulations, we noticed that finer subpopulations of seemingly similar cells within large clusters are often not separated at a satisfactory precision (data not shown). To this end, network based clusters are used as points of reference for further down-sampling of the transcriptomes. DropClust uses an exponential decay function to select higher number of expression profiles from clusters of relatively smaller sizes. Simulated annealing is used to perform hyperparameter search with the aim of restricting the sample size close to a number, manageable by hierarchical clustering. The proposed sampling strategy preserves the rare cell clusters even when the sample sizes are fairly small compared to the population size (**Supplementary Figure 1; Online Methods**).

It is well known that clustering outcome is often improved by careful selection of genes. Principle Component Analysis (PCA) has widely been used for this purpose^7,11^. Traditionally, genes with high loadings on the top few principal components (PCs) are considered to be most informative. This method, in some sense, guarantees selection of the classes of highly variable genes. However, expression variability may not necessarily explain cell type heterogeneity. For gene selection based on high PC loadings, dropClust uses PCs that not only explain a sizable proportion of the observed expression variance but also manifest a large proportion of phenotypic diversity. To this end dropClust uses mixtures of Gaussians to detect PCs with multi-modal distribution of the projected transcriptomes (**Online Methods**). When applied on the real datasets we commonly encountered cases where a top PC featured a small number of modes whereas a trailing PC featured higher levels of modality (**Supplementary Figure 2**). Genes selected in this approach are used for clustering single cell expression profiles using the average linkage hierarchical clustering algorithm. For each of the remaining expression profiles dropClust finds the nearest neighbors from within the sampled transcriptomes. Cluster, that contains the maximum number of neighbors of any transcriptome is assigned as its cluster of origin. **Supplementary Table 1** enlists the the parameters used by the different clustering methods.

Visualizing large-scale scRNA-seq data is challenging. Both Principal Component Analysis (PCA) and t-distributed Stochastic Neighborhood Embedding (tSNE) are widely used used for visualization of scRNA-seq datasets^12^. However, both these methods scale slowly with growing number of transcriptomes. DropClust uses tSNE to obtain the 2D coordinates of a small sub-sample of the data, followed by inferring coordinate pairs of each remaining trancriptome by averaging the coordinates of its nearest neighbors among the sub-sample (**Figure 1A; Supplementary Figure 3, 6; Online Methods**). This strategy offered significant speedup and improved the correspondence between clustering and low dimensional visualization of the data. The complete dropClust work-flow is illustrated in **Supplementary Figure 5**.

**Figure 1.**
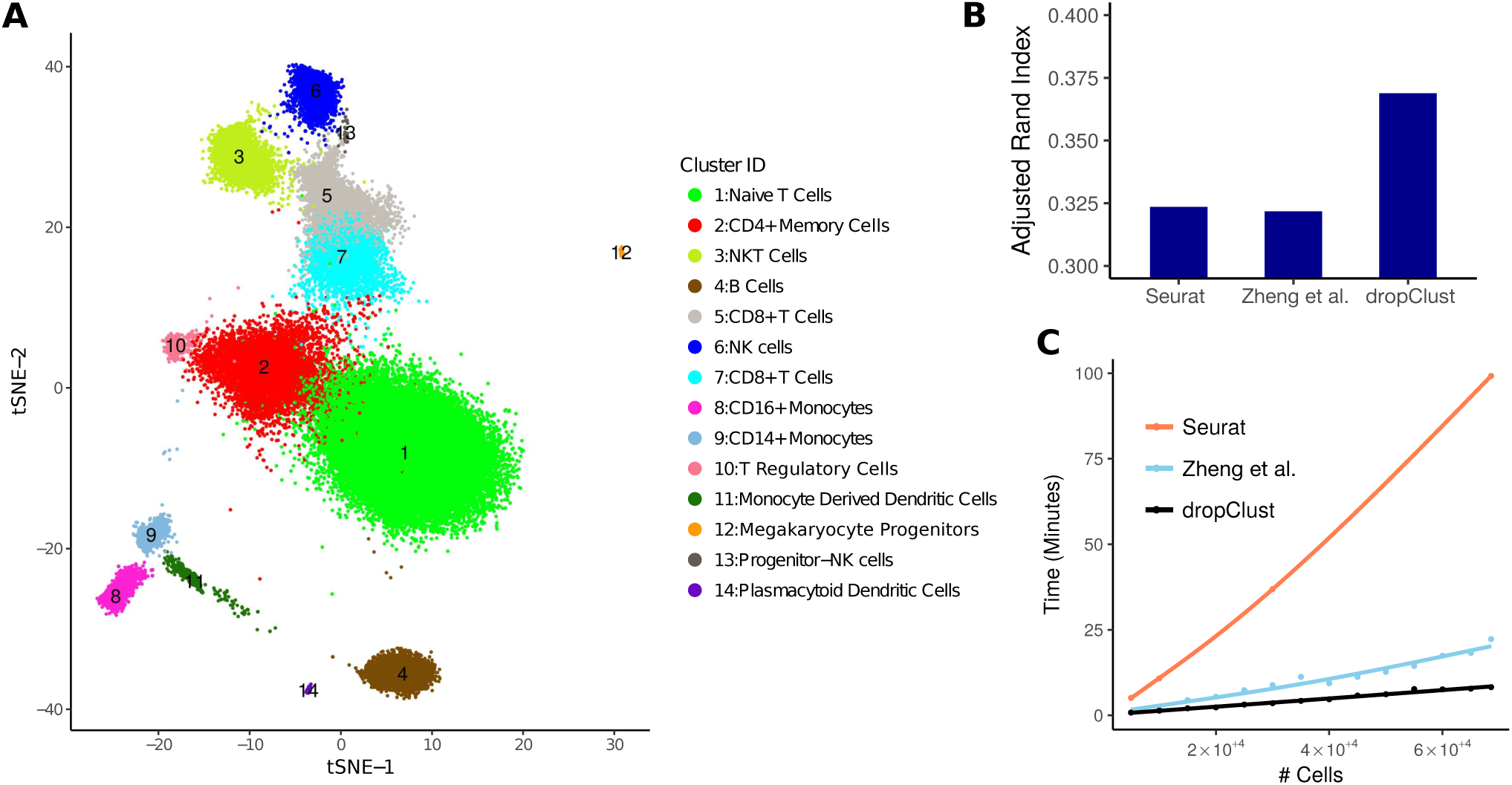
Clustering of ∼ 68k PBMC data. (A) dropClust based visualization (a modified version of tSNE) of the transcriptomes. 14 clusters, retrieved by the algorithm are marked with their respective cluster IDs. Legends show the names of the inferred cell types. (B) Bars show the ARI indexes obtained by comparing clustering outcomes with cell-type annotations. (C) Trend of increase in execution time for different clustering methods with growing number of transcriptomes under analysis.

We applied dropClust first on a collection of ∼ 68k human peripheral blood mononuclear cells (PBMC), annotated based on similarity with matched immune cell subpopulations, purified using Fluorescence-activated cell sorting (FACS)^2^. We identified all major lymphoid and myeloid sub-populations including a number of minor subtypes. Among the populous cell-subtypes we detected naive, memory and cytotoxic T cells, B cells, natural killers (NK), natural kill T cells (NKT cells), CD14+ and CD16+ blood monocytes and monocyte derived dendritic cells. Besides these we also found a number of minor cell types including plasmocytoid dendritic cells, regulatory T cells (Tregs), progenitor NK cells and circulating megakaryocytes progenitors (**Figure 1A**). Differential expression (DE) analysis was carried out between each pair of clusters to identify the cell type specific genes for each sub-population. (**Supplementary Figure 7; Supplementary Table 3**). Details about the mapping of the dropClust predicted clusters to their respective potential cell types can be found in the **Supplementary Material**.

Zheng and colleagues used matched single cell transcriptomes of 11 purified immune cell types for annotating the transcriptomes of the PBMC data^2^. We used this information to benchmark the performance of the cell clustering methods under investigation. For each method, concordance between clusters assignment and cell type annotation was measured by Adjusted Rand Index (ARI). Among all methods, dropClust maximized the ARI (**Figure 1B**).

Besides improved clustering accuracy, dropClust is designed to provide significant speed up. On ∼ 68k PBMC data it took ∼ 8 minutes to perform the clustering. The *k*-means based pipeline proposed by Zheng *et al.* took around 22 minutes whereas it took ∼ 100 minutes for Seurat to generate the clusters. We logged the execution time for different methods while increasing the number of transcriptomes to analyze. Time consumed by dropClust followed a log linear trend w.r.t. cell count. For the other methods time consumption clearly followed non linear growth trajectories (**Figure 1C**).

A major promise of single cell expression profiling at a large scale lies in the possibility of identifying rare cell subpopulations. A cell type may be considered as rare when its abundance in the respective population is ≤ 5%^6,13^. The ability of the clustering methods to detect rare cell-types was assessed through a simulation study. For this, we used a collection of ∼ 3200 scRNA-seq profiles containing Jurkat and 293T cells, mixed *in vitro* at equal proportion^2^. The authors tracked the profile of Single Nucleotide Variants (SNVs) to determine the lineage of the individual cells. The ratio of the two cell types was altered *in silico* by down-sampling one of the populations. Abundance of the minor cell type was varied between 1 to 10 percent. A variant of the popular F1 score was used to measure the algorithm efficacies. DropClust turned out to be the only algorithm that detected the minor clusters nearly accurately at all tested concentrations (**Figure 2; Online Methods**). The existing methods clearly struggled with the smaller concentrations of the rare cell lineage.

**Figure 2.**
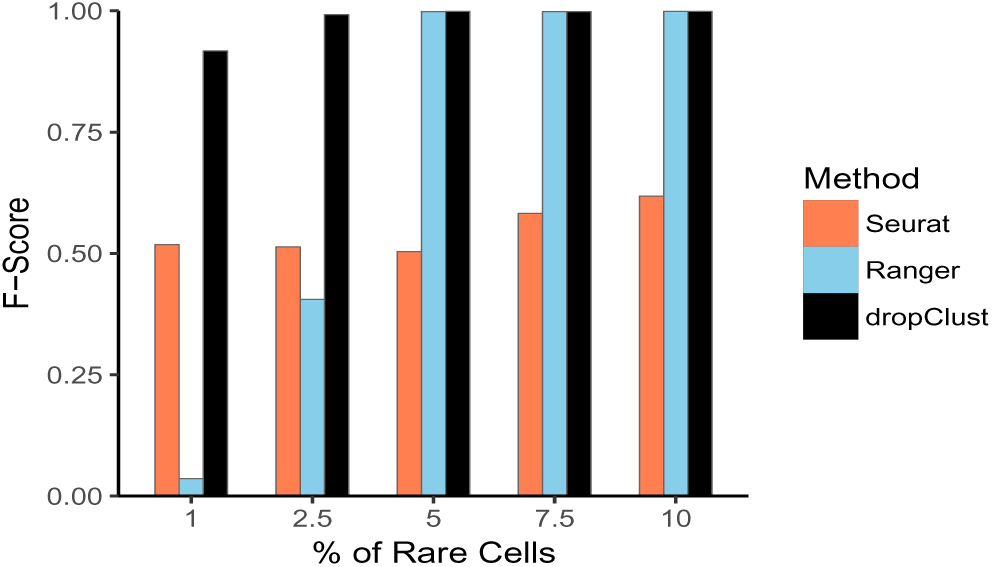
Detectability of minor cell types. Bars showing average of F1-scores, obtained on 10 simulated datasets at each concentration of the minor population. A dataset containing mixture of Jurkat and 293T cells was used for this study.

To rule out the possibility of assay sensitivity, we benchmarked the performance of the clustering methods on two additional Drop-seq datasets from independent studies. The first dataset consisted of ∼ 49k mouse retina cells whereas the second one contained ∼ 2700 mouse embryonic stem cells. Unlike the PBMC data, these two datasets were not supplemented with any secondary source of cell-type identity information. Silhouette Index (SI) was therefore used as an unsupervised measure of clustering accuracy. DropClust yielded the best SI scores on both the datasets, closely followed by the method suggested by Zheng *et. al.* (**Supplementary Material; Supplementary Figure 11**).

The above results clearly demonstrate the superior performance of dropClust in terms of execution time and accuracy of unsupervised cell grouping. Notably, among all tested methods, dropClust demonstrated the maximum sensitivity in detecting the minor cell type. With the increase in single cell transcriptomic throughput capabilities and technology availability (Chromium^*TM*^ by 10x Genomics, ICELL8 by WaferGen Biosystems, similar platform by Illumina and Bio-Rad etc.) we predict unique relevance of the proposed dropClust pipeline.

## Software

The dropClust R package is available at https://github.com/debsin/dropClust.

## Online Methods

### Description of the datasets

For this study we two datasets from a recent work by Zheng and colleagues^2^. The first single-cell-RNAseq (scRNA-seq) data consists of ∼ 68,000 peripheral blood mononuclear cells (PBMC), collected from a healthy donor. Single cell expression profiles of 11 purified subpopulations of PBMCs are used as reference for cell type annotation. This dataset served as a gold standard for performance assessment of the clustering techniques. The second dataset from the same study contains expression profiles of Jurkat and 293T cells, mixed *in vitro* at equal proportions (50:50). All ∼ 3,200 cells of this data are assigned their respective lineages through SNV analysis^2^. Expression matrices for both these datasets were downloaded from www.10xgenomics.com.

We used two additional datasets to benchmark the performance of the clustering algorithms. The first dataset contained transcriptomes of ∼ 49k mouse retina cells^7^ whereas the second data contained transcriptomes of ∼ 2700 mouse embryonic stem cells^14^ (**Supplementary Material**).

### Data preprocessing, normalization and gene selection

Expression matrices for all the datasets were downloaded from publicly available repositories. For each dataset we retained the genes whose UMI counts were > 3 in at least 3 cells. For PBMC data, only ∼ 7,000 genes qualified this criterion. The filtered data matrix was then subjected to UMI normalization that involves dividing UMI counts by the total UMI counts in each cell and multiplying the scaled counts by the median of the total UMI counts across cells^2^. 1000 most variable genes were selected based on their relative dispersion (variance/mean) w.r.t. to the expected dispersion across genes with similar average expression^2,7^.

### Structure preserving sampling of transcriptomes

It is hard to avoid sub-sampling while managing high dimensional genomic data. However, random sub-sampling might result in loss of rare sub-populations. The proposed dropClust pipeline introduces a novel data sampling approach that preserves distinct structural properties of the data. This is achieved in two steps: a) A fairly large (usually minimum of 20000 and a third of the whole population) number of scRNA-seq profiles are randomly selected from the complete set of transcriptomes and then subjected to a fast, approximate graph based clustering algorithm; b) the topological clusters thus obtained are used to guide further sub-sampling of the transcriptomes in a way that retains relatively higher number of cells from smaller clusters, which were otherwise ignored in case of random sub-sampling.

To construct the network, top-*k* approximate nearest neighbors (*k*=10 by default) are identified rapidly by employing Locality Sensitive Hashing (LSH)^15^. A faster and more accurate implementation of the original LSH, called LSHForest is used for this purpose^16^. The nearest neighbor network (NNN) of transcriptomes thus created is subjected to Louvain^10^, a widely used method for detecting community structures in networks. Notably, Seurat^7^ uses Shared Nearest Neighbors (SNN) for network construction at one specific stage. While construction of NNN using LSH takes *O*(log *n*) time^16^, building SNN requires *O*(*n*^2^) time^8^, where *n* denotes the number of single cell expression profiles. The choice of LSH to search nearest neighbors leads to a dramatic reduction in computation time. Since LSH is an approximate method for nearest neighbor search, the clusters obtained need further refinement. Moreover, Louvain offers limited control for determining cluster resolution.

To ensure selection of sufficient representative transcriptomes from small clusters, an exponential decay function^17^ is used to determine the proportion of transciptomes to be sampled from each cluster. For *i^th^* cluster, the proportion of expression profiles *p_i_* was obtained as follows.

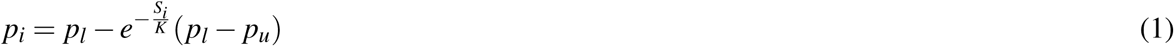

where *S_i_* is the size of cluster *i*, *K* is a scaling factor, *p_i_* is the proportion of cells to be sampled from the *i^th^* Louvain cluster. *p_l_* and *p_u_* are lower and upper bounds of the proportion value respectively. Based on the above equation we may show the following:

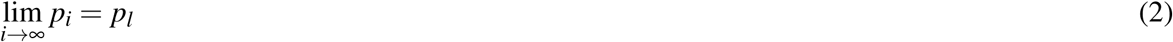

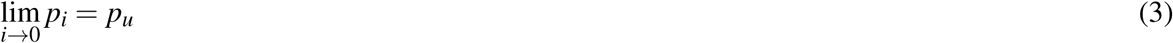

Since Equation 1 does not explicitly impose any upper bound on the final sample size, one may be left with an arbitrarily high or low number of single cell transcriptomes for final clustering. To address this, dropClust allows user specify his preferred sample size and employs simulated annealing (SA)^18^ to come up with the right values for *p_l_, p_u_* and *K*. This operation may formally be described as follows.

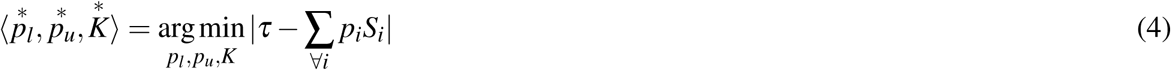

where *τ* denotes the user specified sample size. We used simulated annealing implementation from the `GenSA` R package^19^.

### Clustering of sampled cells

For the cells obtained through structure preserving sampling, gene selection is performed based on Principal Component Analysis (PCA). For each of the top 50 Principal Components (PC), we estimate the explained heterogeneity by inspecting the multi-modal nature of its marginal distribution. Gaussian Mixture Model (GMM)^20^, supplemented with Bayesian Information Criterion (BIC)^21^ is used to determine the number of modes corresponding to each PC. Each of these modes is expected to represent a cell type. R package `mclust` is used for this purpose. PCs, modeled by three or more Gaussians are used for PC-loading based gene selection^11^. Top 200 high-loading genes are retained for the subsequent clustering step.

Average-linkage hierarchical clustering is performed to group the sampled cells based on expression of the 200 selected genes. Euclidean distance is used as the measure of dissimilarity. To cut the dendrogram `cutreeDynamic()` is used from the `dynamicTreeCut` R package^22,23^.

### Post-hoc cluster assignment for left out transcriptomes

Cells that are not subjected to hierarchical clustering are assigned their respective clusters of origin using a simple post-hoc cluster assignment strategy. To achieve this, locality preserving hash codes are generated for the clustered transcriptomes, using `LSH-Forest`. For each of the left out transcriptomes *k* (*k* = 5, by default) approximate nearest neighbors are then found through LSH queries. Each unallocated transcriptome is assigned the cluster of origin for which the most number of representatives are found in its corresponding set of *k* nearest neighbors. Ties for cluster assignment are broken at random.

### 2D embedding of transcriptomes for visualization

The 2D embedding of samples is carried out in two steps. In the first step t-SNE is applied to transcriptomes obtained through structure preserving sampling. Top 200 PCA-selected genes are used for this purpose. In the next step, remaining transcriptomes are allocated positions in the pre-existing 2D map of the sampled cells. To perform this, we borrow the sets of *k* nearest neighbors, found at the time of post-hoc cluster assignment. Coordinates for each newly added point are derived by averaging t-SNE coordinate values of neighbors that belonged to its cluster of origin.

### Differential Expression of Genes

To speed up the differential expression (DE) analyses, we consider 100 randomly chosen transcriptomes from each cluster. Only genes with count > 3 in at least 0.5% of these cells are retained for the analysis. Fast nonparametric, DE analysis tool NODES is used to to make DE gene calls with 0.05 as the cut off value for false discovery rate (FDR) and a fold change of 1.2^24^. Among the DE genes, ones that are significantly upregulated in a specific cluster w.r.t. each of remaining clusters are named cell type specific genes.

### Rare sub-population detectability

The dataset containing Jurkat and 293T cells at equal ratio was used for performing simulations to assess detectability of minor cell populations. Cell type identity of each transcriptome of this dataset was determined by SNV analysis^2^. To introduce rareness, we forcibly reduced the frequency of one of the cell types. To prevent bias, we performed these experiments by treating both cell types as rare in separate simulations. In simulated datasets, the proportion of rare cell transcriptomes was varied among 1%, 2.5%, 5% and 10%. For each of these specified concentrations, 10 datasets were created by independent sub-sampling of transcriptomes of a specific type. The transcriptomes of the major cell type were not subjected to any kind of sampling. Since this procedure was repeated for both the cell types, for each concentration a total of 20 datasets were produced.

We used F1-score as a measure for detectability of rare cell clusters. The score is defined as follows.

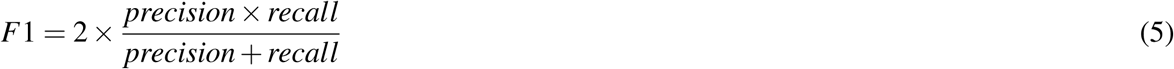

To compute the above score we first associated the the predicted cluster that contained the majority of the rare cells to the rare cell group. Following this, *recall* was defined as the ratio between the number of true rare cells within the predicted rare cell cluster and the total number of known rare cells. On the other hand, *precision* was defined as the ratio between the number of known rare cells within the predicted rare cell group and the total number of cells in the predicted rare cell group.

## Supplementary Material

**Supplementary Figure 1.**
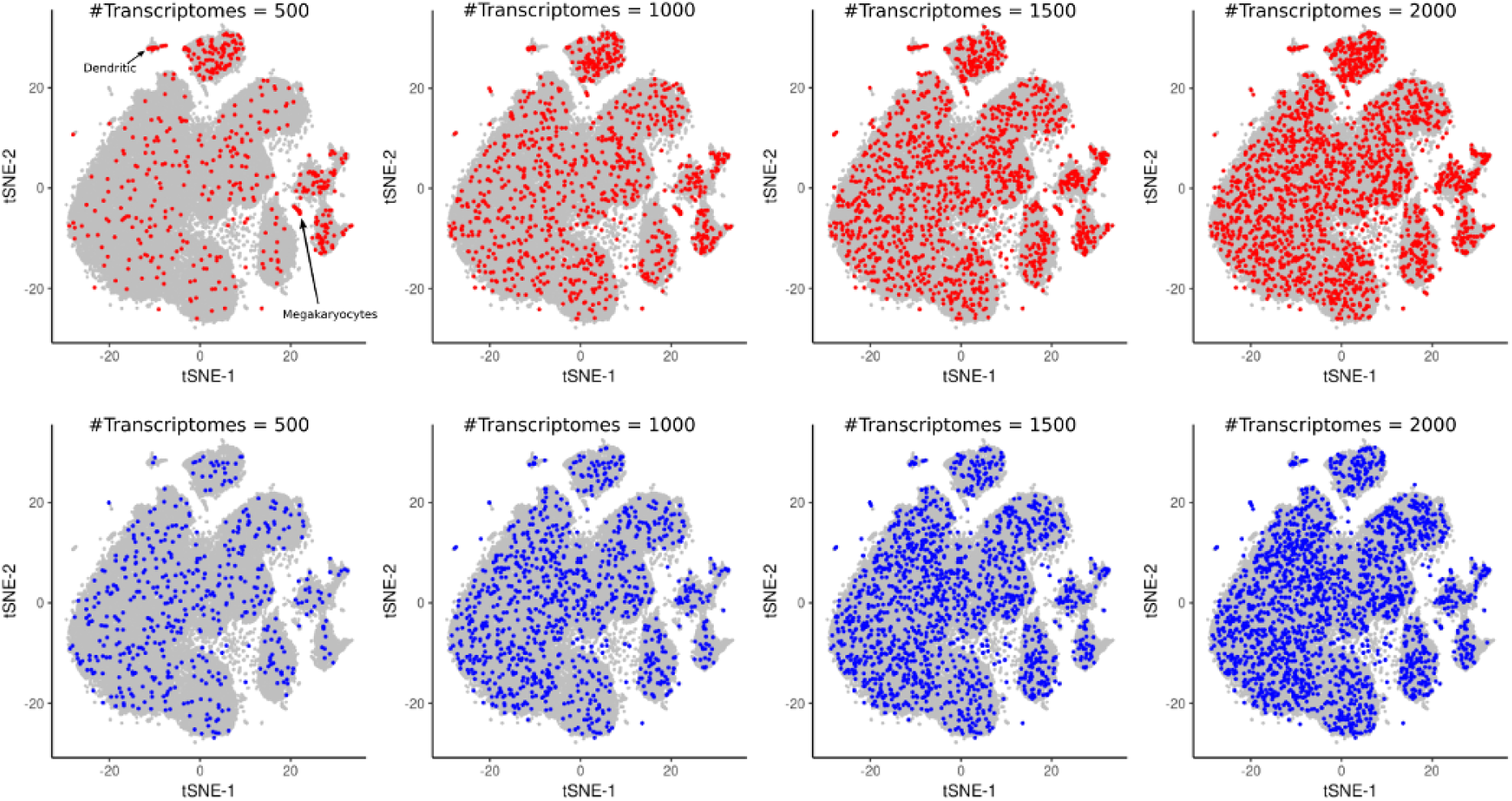
Comparison between structure preserving sampling and random sampling. The former method retains more representative transcriptomes from less prevalent cell types when sample size is small. To see the impact of structure preserving sampling, sample size was varied between 500 to 2000. Red dots represent transcriptomes selected by the structure preserving sampling method whereas blue dots are selected by random sampling. Coordinates for 2D embedding are sourced from Zheng *et al.*

**Supplementary Figure 2.**
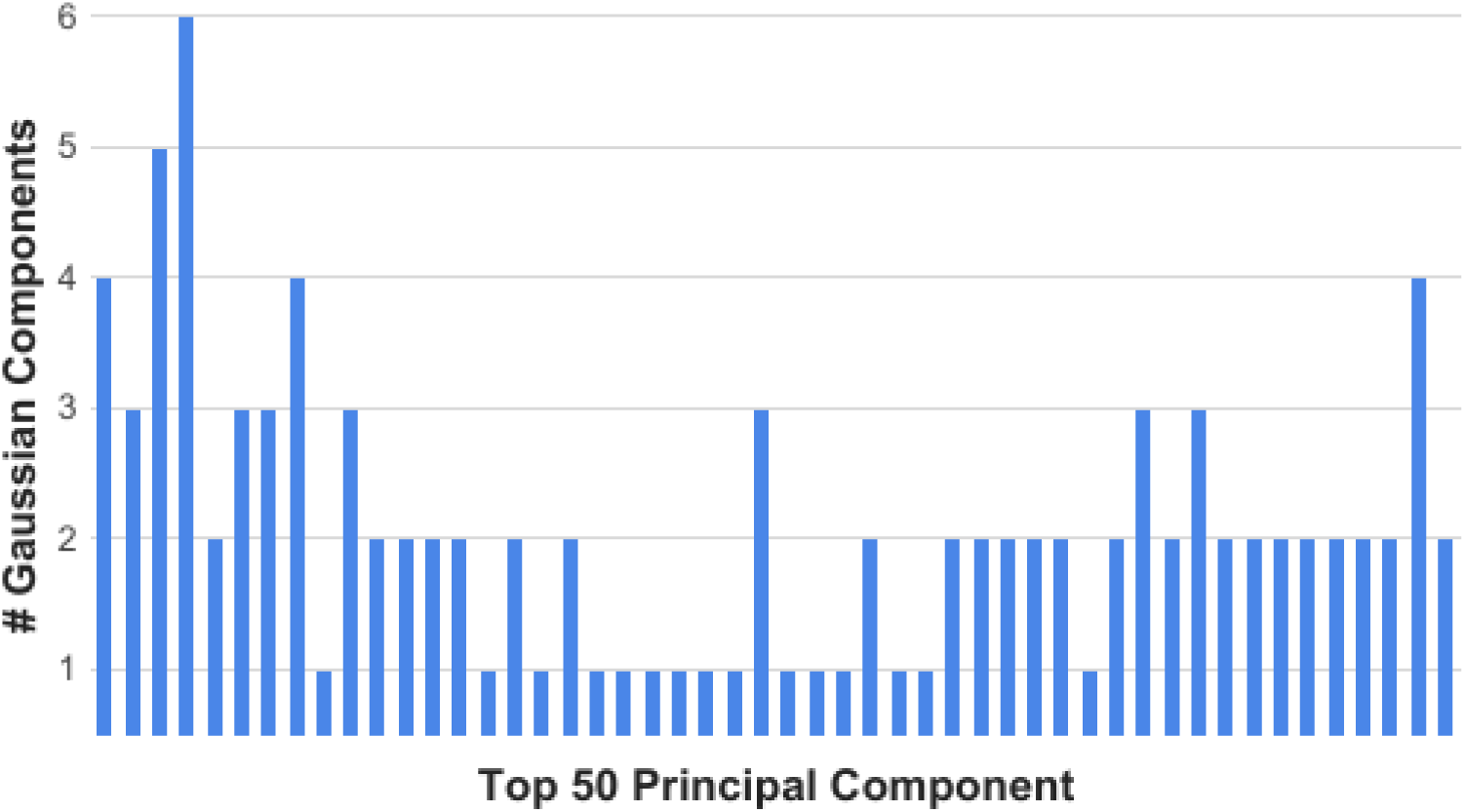
Barplot depicting the number of estimated Gaussian components for each of the top 50 principal components derived from the PBMC data.

**Supplementary Table 1.** Details of clustering parameters used by each method for the respective datasets.

| **Dataset** | **Method** | **Clustering Algorithm** | **Parameter Description** |
| --- | --- | --- | --- |
| PBMC | dropClust | Hierarchal Clustering | minClusterSize = 20, deepSplit = 3 |
| PBMC | Zheng *et al.* | K-Means | k = 10 |
| PBMC | Seurat | SNN | resolution = 0.6 |
| Jurkat:293T | dropClust | Hierarchal Clustering | minClusterSize = 80% of smaller cluster, deepSplit = 0 |
| Jurkat:293T | Zheng *et al.* | K-Means | k = 2 |
| Jurkat:293T | Seurat | SNN | resolution = 0.6 |
| Retina | dropClust | Hierarchal Clustering | minClusterSize = 22, deepSplit = 1 |
| Retina | Zheng *et al.* | K-Means | k = 20 |
| Retina | Seurat | SNN | resolution = 0.5 |
| ESC | dropClust | Hierarchal Clustering | minClusterSize = 20, deepSplit = 0 |
| ESC | Zheng *et al.* | K-Means | k = 4 |
| ESC | Seurat | SNN | resolution = 0.4 |

**Supplementary Figure 3.**
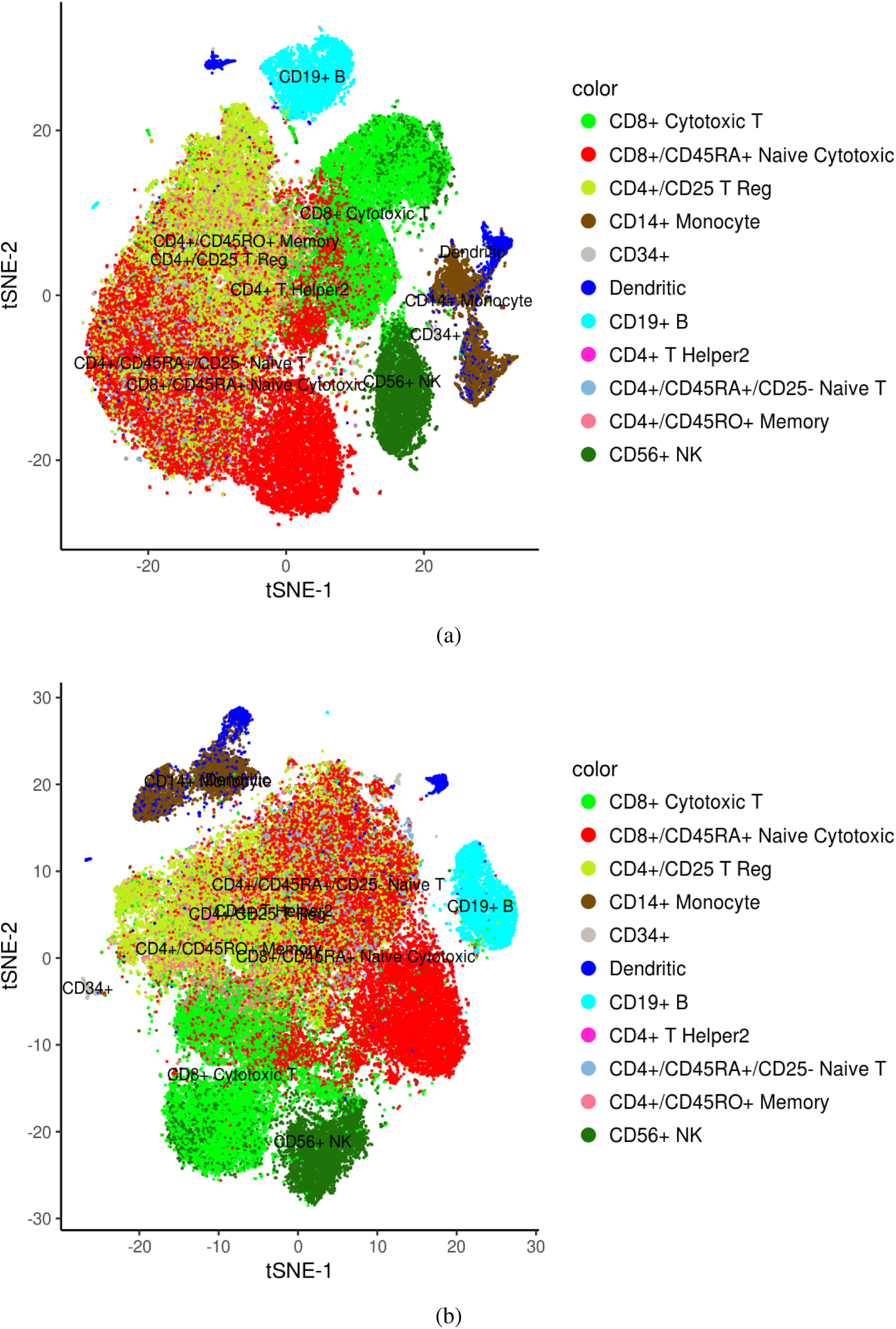
(a) 2D embedding of 68K PBMCs using the methodology proposed by Zheng *et al.*^1^; (b) Similar 2D embedding using Seurat^2^. Note: Cells are colored as per the cell type annotations provided by Zheng *et al.*^1^

**Supplementary Figure 4.**
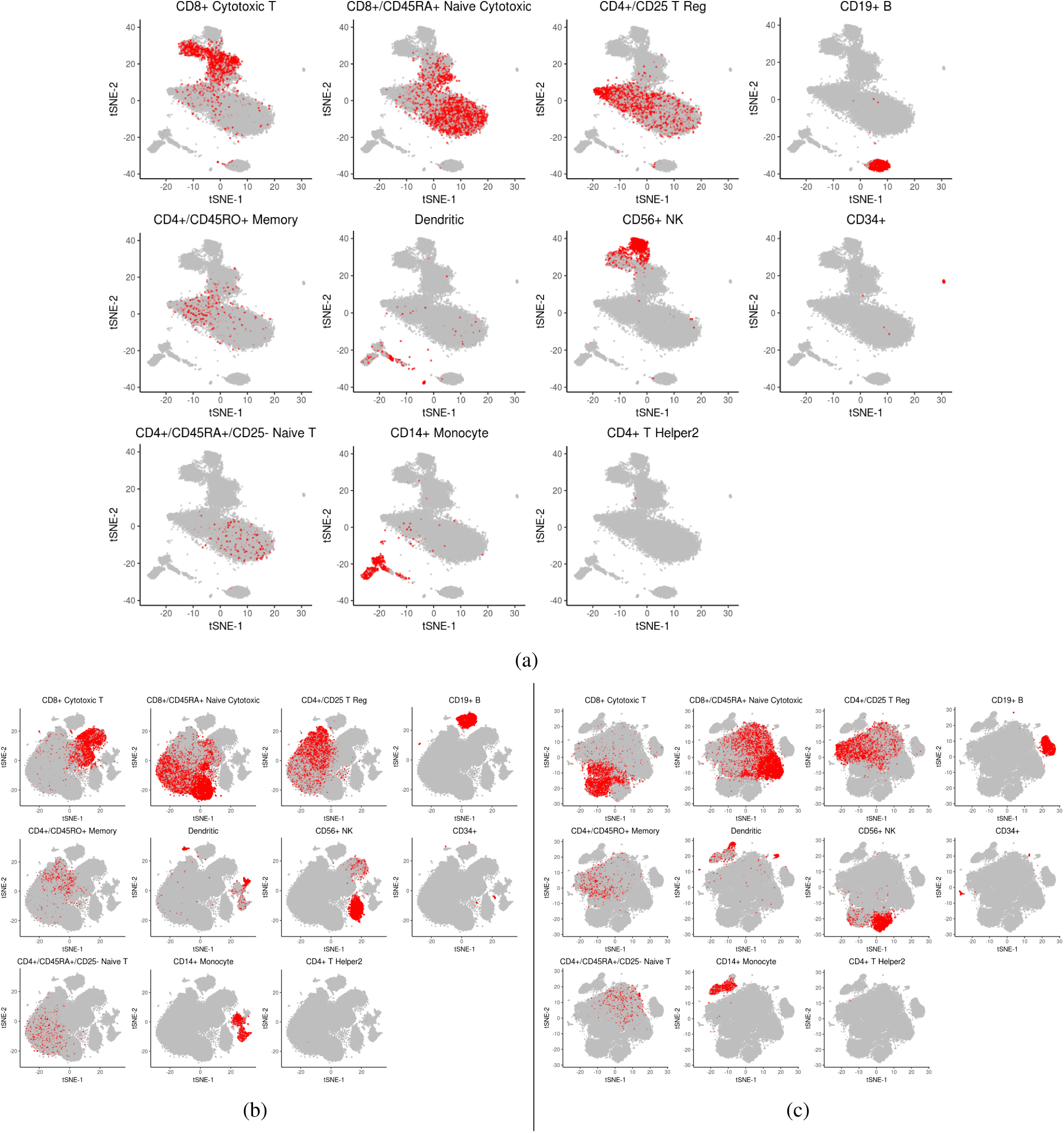
(a) Localization of PBMC transcriptomes of same type (based on annotation) on the 2D embedding produced by dropClust. (b) Similar plots with background 2D map generated using the methodology proposed by Zheng et al.^1^ (c) Similar plots for Seurat^2^.

**Supplementary Figure 5.**
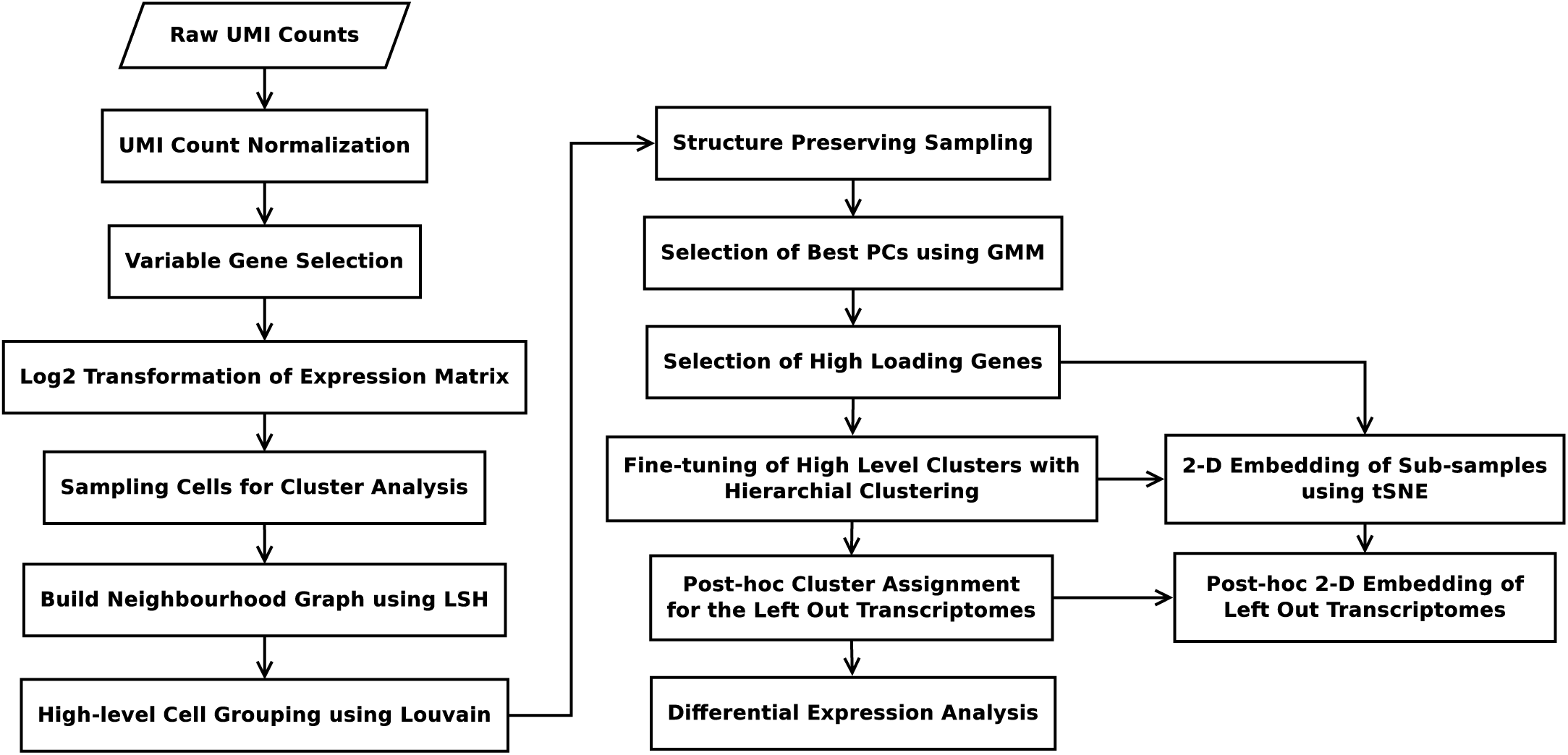
Flowchart of dropClust pipeline.

**Supplementary Figure 6.**
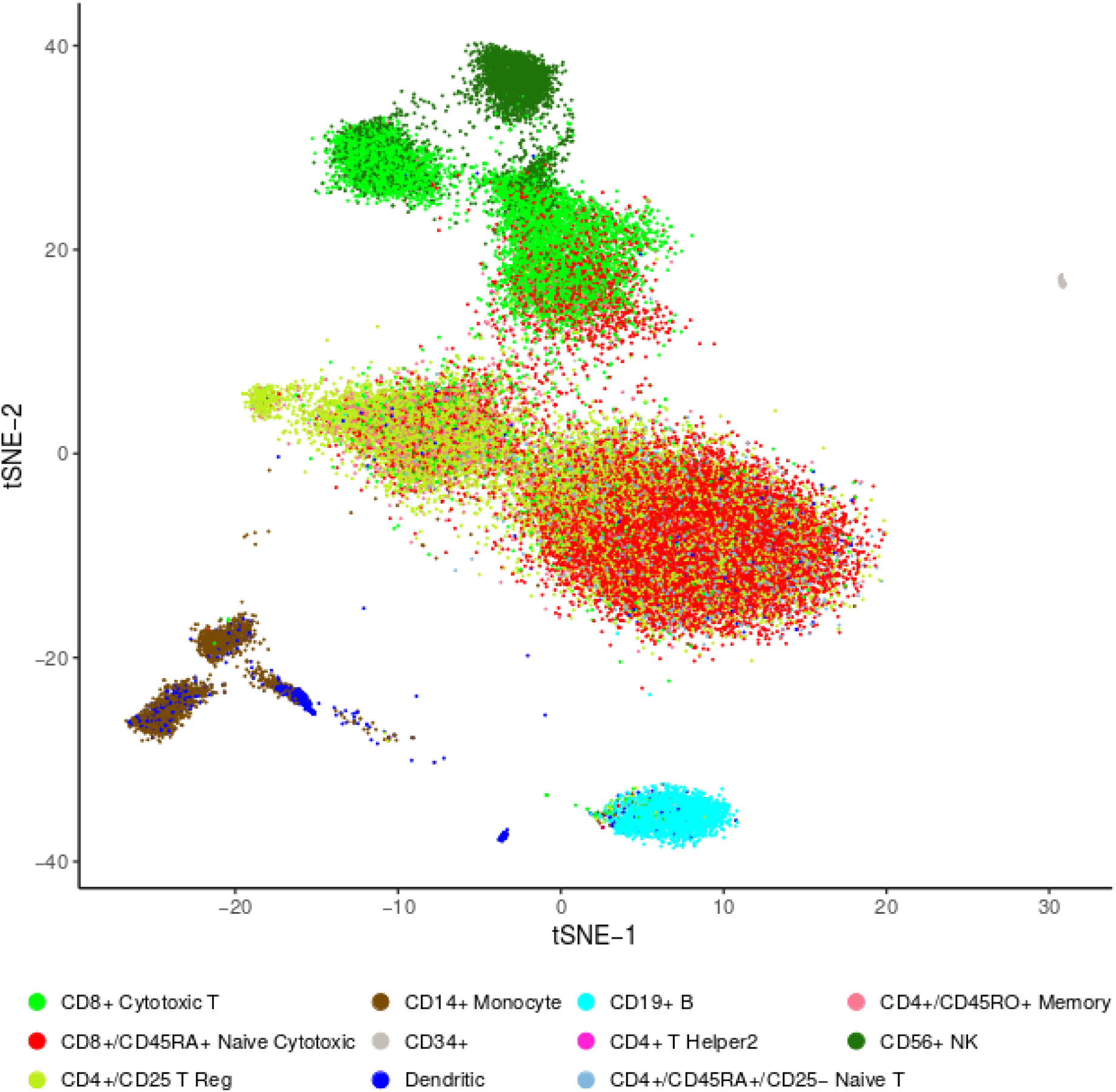
2D embedding of 68K PBMCs using dropClust. The cells are colored as per the cell type annotations provided by Zheng *et al.*^1^

**Supplementary Figure 7.**
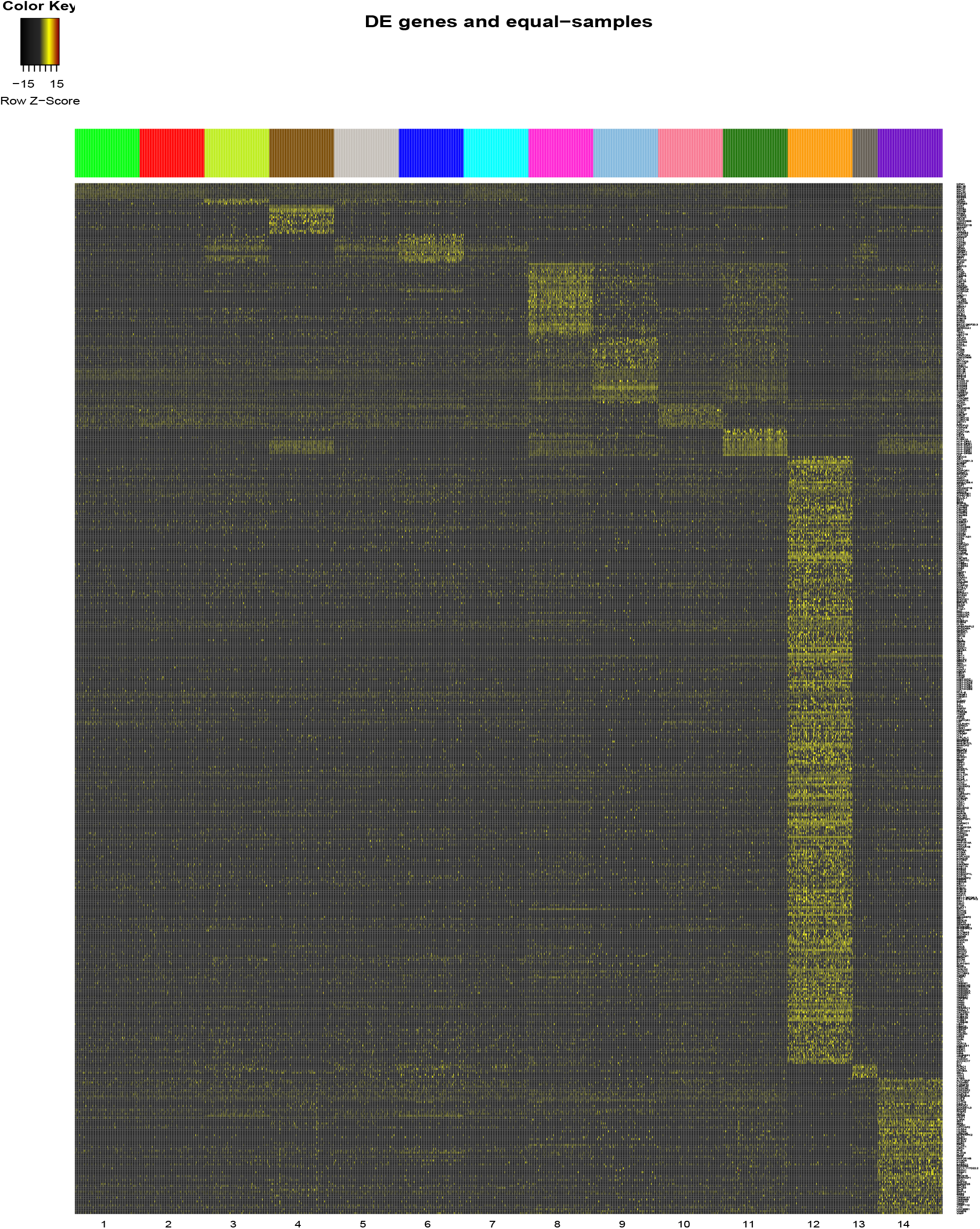
Heatmap of cell type specific differentially up-regulated genes. No such genes were found for Clusters 2, 5 and 7.

## 1 Cell type determination

We could associate the predicted groups of transcriptomes with known cell-types based on marker gene expression. Such associations were not always unambiguous. There are two principal reasons for such ambiguities: 1. Surface protein concentration is not always linearly related to the expression of the corresponding gene. Well known surface markers are commonly found having low expression. 2. High drop-out rates and lack of sequencing depth cause prevalence of zeros as expression estimate. As a result, cell type specific yet low expressed genes are often not detected in single cell assays. Under these constraints, we tried to gather as much evidence as possible to assign a putative cell type to each of the detected PBMC clusters. Supplementary Figure 7 shows the heatmap of the cell type specific differentially up-regulated genes.

**Cluster 1 (*% of Cells*: 46) *Predicted cell type***: Naive T cells. ***Evidence***: 1. Based on transcriptomic similarity with the purified PBMC subpopulations (see Supplementary Figure 4 and Supplementary Figure 8), a majority of the CD8+ and CD4+ naive T cells are present in this cluster. 2. High expression levels of CCR7 and CD27 indicate that many of the cells present in this cluster are either CD4+ naive or CD4+ memory T cells. However, as per the annotations (based on transcriptomic similarity, see Supplementary Figure 8) CD4+ naive T cells are way more enriched in this cluster than the memory cells. 3. High expression levels of CD8A and CD8B, which indicate the presence of CD8+ T cells (Supplementary Figures 9 & 10).
**Cluster 2 (*% of Cells*: 14.9) *Predicted cell type***: CD4+ Memory cells. ***Evidence***: 1. Based on transcriptomic similarity with the purified PBMC subpopulations (see Supplementary Figure 4 and Supplementary Figure 8), the majority of the memory T cells are present in this cluster. 2. IL7R (CD127) and CD27 are well expressed in this population^3,4^ whereas the average expression of CCR7 is low as compared to its expression in cluster 1 (Naive T cells). 3. A fraction of the CD4+ regulatory T cells (as per annotation) are present in this cluster, indicating that the cluster mostly harbors cells of various CD4+ subtypes. Treg annotated cells, present in Cluster 2 were indistinguishable from the memory T cells. An explanation for that may stem from the fact that both CD4+ T cells and Tregs originate from thymus^5^. Of note, it is possible that some of the memory cells in this cluster were mis-annotated as T cells by Zheng *et al*^1^.
**Cluster 3 (*% of Cells*: 9) *Predicted cell type***: Natural Killer T (NKT) cells. ***Evidence***: 1.Based on annotations, cluster 3 shares a large fraction of cytotoxic T cells and a tiny fraction of NK cells (see Supplementary Figure 4 and Supplementary Figure 8). 2. ZNF683, a known NKT cell marker is found to be exclusively up-regulated in this group^6,7^. 3. CD8A and CD8B are well expressed in this cluster, indicating its proximity to CD8+ T cells population. 4. A number of Natural Killer markers including NKG7, GNLY, GZMB are up-regulated in this cluster (Supplementary Figure 9).
**Cluster 4 (*% of Cells*: 5.8) *Predicted cell type***: B cells. ***Evidence***: 1. Canonical B cell markers CD79A and CD37 are exclusively highly expressed in this cluster (Supplementary Figure 9). 2. As per annotations almost all B cells are localized in this cluster (Supplementary Figure 4 and Supplementary Figure 8).
**Cluster 5 & 7 (*% of Cells*: 11.8) *Predicted cell type***: CD8+ T cell subtypes. ***Evidence***: 1. Based on annotations, cells in these clusters are most similar to purified CD8+ T cell subpopulations (see Supplementary Figure 4 and Supplementary Figure 8). 2. CD8A and CD8B are well expressed in this cluster. 3. Cell type specific genes were not found for these clusters except GZMK for cluster 7. GZMK is known to have differential expression patterns across NK and CD8+ T cell subtypes^8^.
**Cluster 6 (*% of Cells*: 4.9) *Predicted cell type***: Natural Killers. ***Evidence***: 1. As per the annotations a majority of the NK cells are localized in this cluster (Supplementary Figure 4 and Supplementary Figure 8). 2. A number of well-known NK cell markers including CD160^9,10^, NKG7^11^, GNLY^12^, CD247^13^, CCL3^14^ and GZMB^15^ were found to be differentially up regulated (Supplementary Figure 9) in this group.
**Cluster 8 & 9 (*% of Cells*: 4.9) *Predicted cell type***: CD16+ and CD14+ Monocytes respectively. ***Evidence***: 1. The majority of annotated monocytes are localized in these clusters (Supplementary Figure 4 and Supplementary Figure 8). 2. The overall high expression of CD16^16^ and CD68^17^ in Cluster 8 (Supplementary Figure 9) indicates that the cluster indeed represents the CD16+ Monocyte sub-population. 3. On the other hand, the overall high expression of CD14^16^ and S100A12^18,19^ in Cluster 9 (Supplementary Figure 9) indicates that the cluster most likely represents the CD14+ Monocyte sub-population.
**Cluster 10 (*% of Cells*: 0.48) *Predicted cell type***: Regulatory T (Treg) cells. ***Evidence***: 1. The majority of the cells in this cluster match with the purified Regulatory T cell subpopulation (Supplementary Figure 4 and Supplementary Figure 8). 2. Among the Treg cell markers CD52, CCR10, CMTM7^20^ (Supplementary Figure 9) were found to be highly expressed. FOXP3 and CD25 were also expressed at higher levels (Supplementary Figure 10).
**Cluster 11 (*% of Cells*: 1.5) *Predicted cell type***: Monocyte Derived Dendritic Cells. ***Evidence***: 1. Based on the annotations, this cluster mostly consists of Dendritic cells (Supplementary Figure 8). 2. A fraction of the transcriptomes of this cluster is annotated match purified monocyte subpopulation (Supplementary Figure 4 and Supplementary Figure 8). 3. An overall high expression of CST3, a Monocyte marker, which is also known to be highly expressed in Monocyte Derived Dendritic cell population^21^. 4. Other Monocyte Derived Dendritic cell markers including CD1C^22,23^ and FCER1A^21^ were found differentially upregulated in this cluster (See Supplementary Figure 9).
**Cluster 12 (*% of Cells*: 0.2) *Predicted cell type***: Circulating Megakaryocyte Progenitors. ***Evidence***: 1. Differential up-regulation of Megakaryocyte markers PF4^24^, PPBP^25^ and PLA2G12A^26^ (Supplementary Figure 9). 2. Transcriptomes in this cluster match strikingly with purified CD34+ population (Supplementary Figure 4 and Supplementary Figure 8). It is a well known fact that Megakaryocyte progenitors express CD34 antigen^27^.
**Cluster 13 (*% of Cells*: 0.1) *Predicted cell type***: Natural Killer Progenitors (NKP) ***Evidence***: 1. Differential up-regulation of ID2 (Supplementary Figure 9), an indicator of commitment to NK cells^28,29^. 2. Overall high expression of NK cell specific markers - GNLY and NKG7 (Supplementary Figure 9).
**Cluster 14 (*% of Cells*: 0.3) *Predicted cell type***: Plasmacytoid Dendritic Cells. ***Evidence***: 1. Up-regulation of GZMB (Supplementary Figure 9), which is known to be highly expressed in both NK cells and Plasmacytoid Dendritic cells is both a marker of NK cells and Dendritic cells^30^. 2. Differential up regulation of well known Plasmacytoid Dendritic cell marker CD123 (IL3RA)^31^ (Supplementary Figure 9). 3. As per the cell type annotation, a majority of the transcriptomes of this cluster match with purified Dendritic cell population (Supplementary Figure 8).

Some of the well-known markers like CD4 or CD8B which either failed to exhibit any cell type specific up-regulation or qualify the gene selection criteria are provided in Supplementary Figure 10 for reference. The marker genes used in the cell-type determination are listed in Supplementary Table 2. The list of cell type specific genes for each cluster are mentioned in Supplementary Table 3.

**Supplementary Table 2.** Markers used for cell type inference.

| Cluster ID | Potential cell type | Markers |
| --- | --- | --- |
| 1 | Naive T Cells | CD27<sup>32</sup>, CCR7<sup>33</sup>, CD8A , CD8B* |
| 2 | CD4+ Memory Cells | IL7R<sup>34</sup>, CD27<sup>32</sup>, CCR7<sup>33</sup> |
| 3 | **NKT Cells** | ZNF683<sup>6,35</sup> (UniProtKB - Q8IZ20), CD8A , CD8B* |
| 4 | B Cells | CD79A<sup>7</sup>, CD37<sup>36</sup>. |
| 5 & 7 | CD8+ T Cells | GZMK<sup>8</sup>, CD8A , CD8B* |
| 6 | NK cells | CD160<sup>9,10</sup>, NKG7<sup>11</sup>, GNLY<sup>12</sup>, CD247<sup>13</sup>, CCL3<sup>14</sup>, GZMB<sup>15</sup> |
| 8 & 9 | CD16+ and CD14+ Monocytes | CD68<sup>17</sup>, CD16 (FCGR3A)<sup>16</sup>, CD14<sup>16</sup>, S100A12<sup>18,19</sup> |
| 10 | **Regulatory T Cells** | CCR10, CD25(IL2RA)*, CD52<sup>37</sup>, CMTM7, FOXP3<sup>20*</sup> |
| 11 | Monocyte Derived Dendritic Cells | CST3<sup>21</sup>, CD1C<sup>22,23</sup>, FCER1A<sup>21</sup> |
| 12 | Megakaryocyte Progenitors | PF4<sup>24</sup>, PPBP<sup>25</sup>, PLA2G12A<sup>26</sup> |
| 13 | **Progenitor-NK cells** | ID2<sup>28,29</sup> |
| 14 | **Plasmacytoid Dendritic Cells** | GZMB<sup>30</sup>, CD123 (IL3RA)<sup>31</sup> |

\* Markers that are well-known but failed to qualify the gene selection criteria.

**Supplementary Figure 8.**
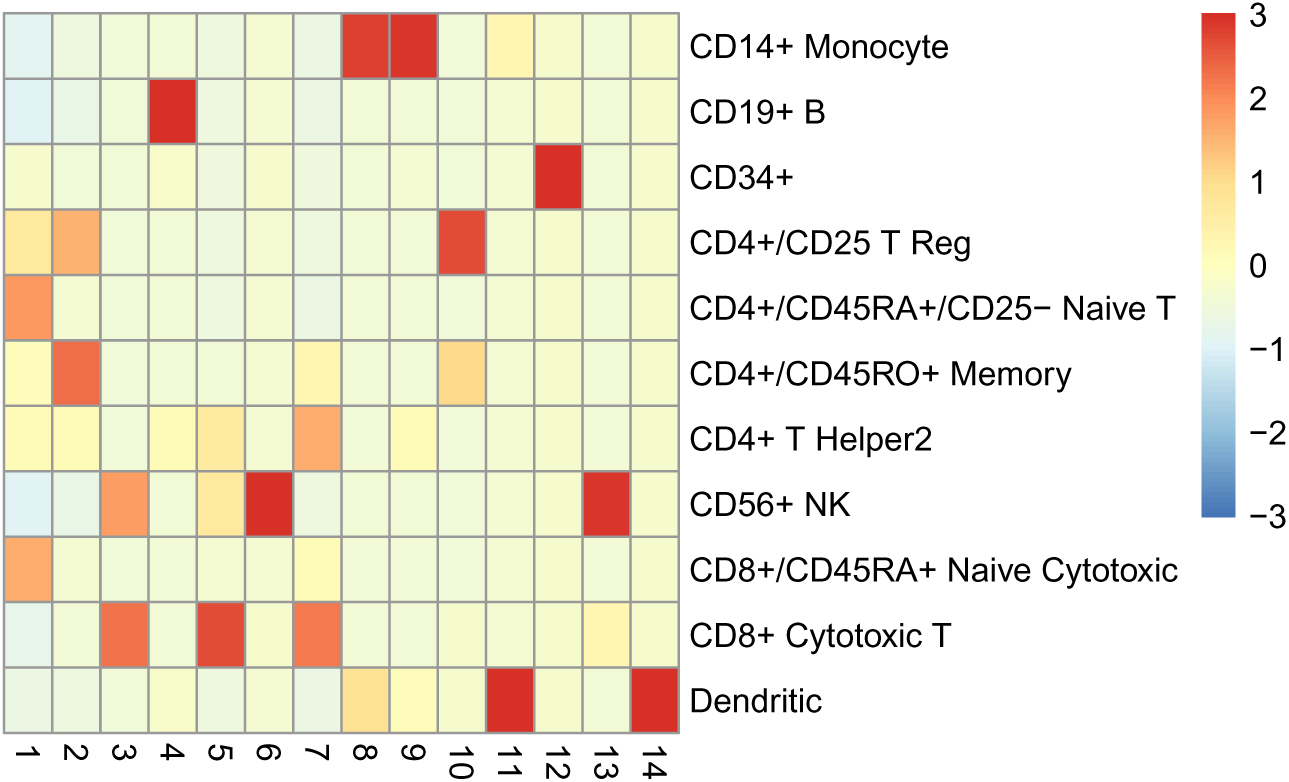
Relative enrichment of annotated cell types in the clusters obtained from the PBMC data.

**Supplementary Figure 9.**
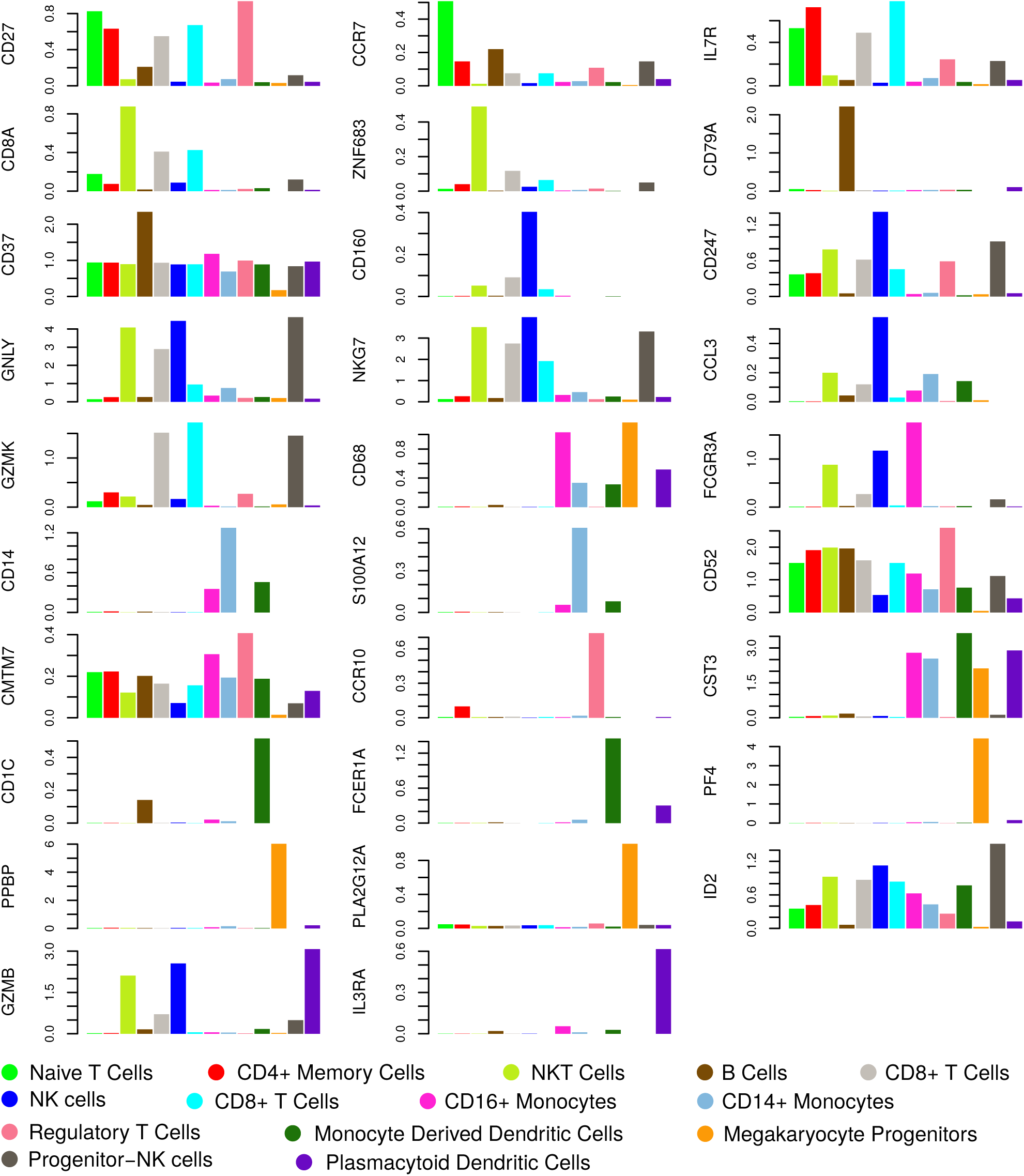
log2 of average expression of the marker genes across the predicted cellular sub-populations (clusters).

**Supplementary Figure 10.**
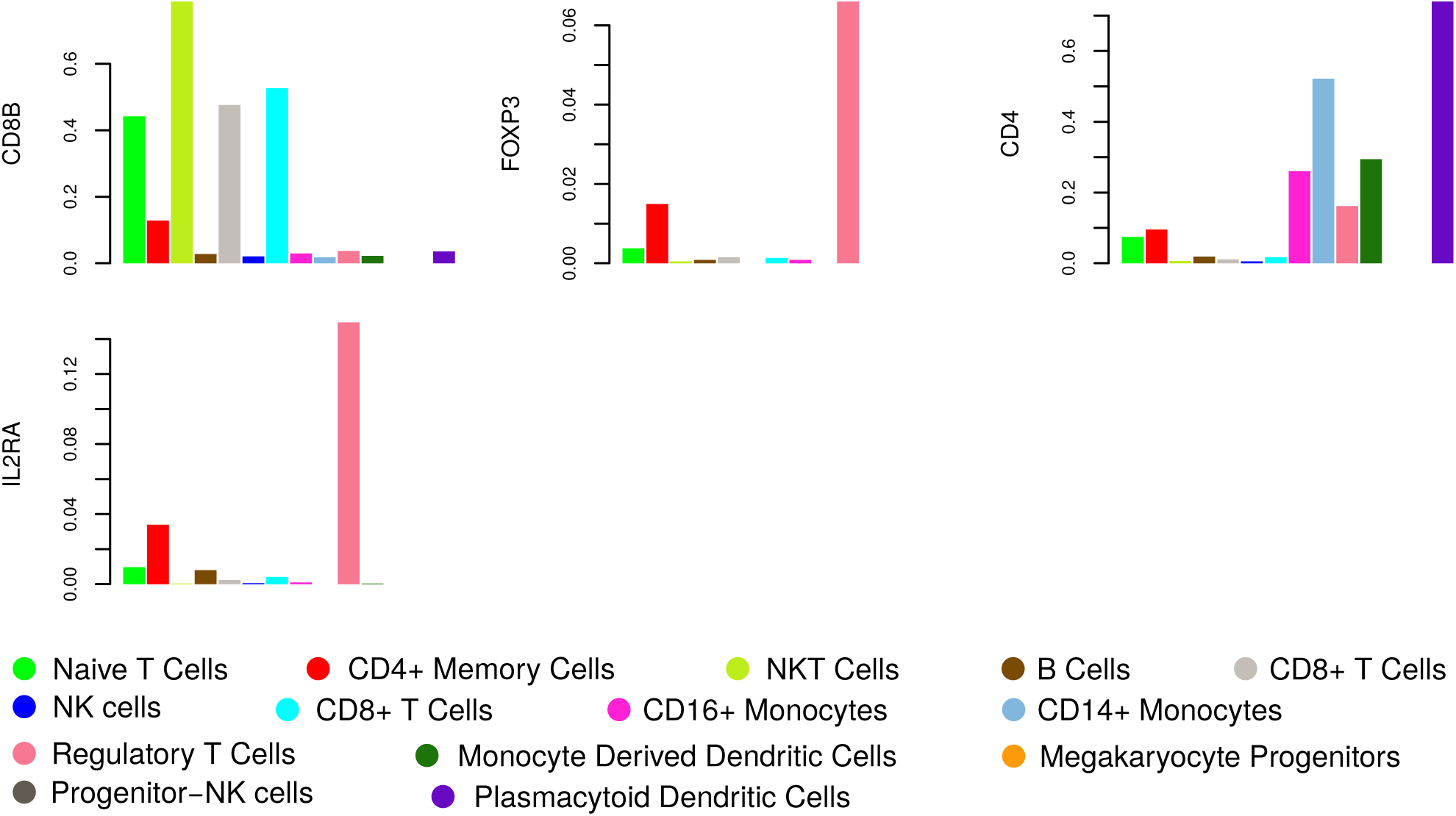
log2 of average expression of the markers that are well-known but failed to qualify the gene selection criteria.

## 2 Results on additional datasets

We evaluated dropClust on two additional datasets. The first dataset consists of transcriptomes of 49,300 mouse retina cells (`GSE63473`)^2^ and the second dataset contains expression profiles of ∼ 2700 mouse embryonic stem cells (ESC) (`GSE65525`)^38^. Both the studies are exploratory in nature and therefore lack any secondary source of information for lineage determination. For these datasets, we, therefore, computed the Silhouette scores (a popular unsupervised metric of cluster quality) corresponding to the cell groupings obtained using different clustering methods. Silhouette is a non parametric measure of the trade off between cluster tightness and inter-cluster separation^39^. For large sample sizes, it takes a long time to compute Silhouette score. To this end, we created 100 independent sets of 500 transcriptomes through bootstrapping. Average Silhouette scores thus obtained are depicted through the boxplots in Supplementary Figure 11. Parameter values used by different clustering methods are furnished in Supplementary Table 1.

**Supplementary Figure 11.**
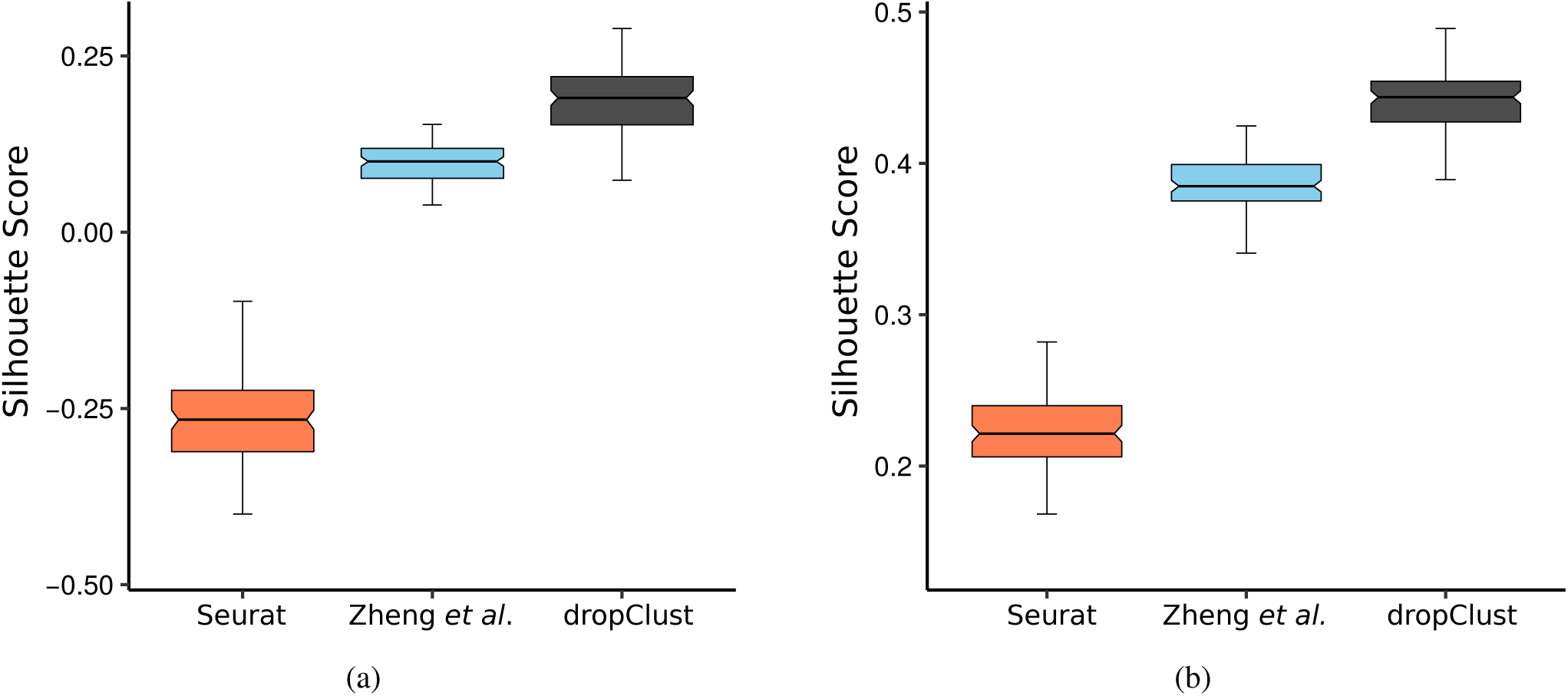
(a) Boxplots depicting average Silhouette scores computed on 100 bootstrap samples from the mouse retina cell data^2^. A separate boxplot is used for each concerned clustering method. (b) Similar plots for the mouse ESC dataset^38^.

